# Development of a robust automated tool for the annotation of embryo morphokinetic parameters

**DOI:** 10.1101/445288

**Authors:** M Feyeux, A Reignier, M Mocaer, J Lammers, D Meistermann, S Vandormael-Pournin, M Cohen-Tannoudji, P Barrière, P Barrière, P Paul-Gilloteaux, L David, T Fréour

**Affiliations:** CRTI UMR1064, INSERM, Université de Nantes, Nantes, France; ITUN, CHU Nantes, Nantes, France; Service de médecine et biologie du développement et de la reproduction, CHU Nantes, Nantes, France; Faculté de médecine, Université de Nantes, Nantes, France; SFR Santé, MicroPICell core facility, CNRS, INSERM, Université de Nantes, Nantes, France; Early Mammalian Development and Stem Cell Biology, Department of Developmental & Stem Cell Biology, Institut Pasteur, 25 rue du Dr. Roux, 75015, Paris, France; CNRS UMR 3738, Institut Pasteur, Paris 75015, France; SFR-SANTE, iPSC core facility, CNRS, INSERM, Université de Nantes, Nantes, France

**Keywords:** IVF, time-lapse imaging, automation

## Abstract

**Study Question:** Is it possible to automatically annotate human embryo development in time-lapse devices, with results comparable to manual annotation?

**Summary Answer:** We developed an automated tool for the annotation of embryo morphokinetic parameters having a high concordance with expert manual annotation in a large scale-study.

**What is Known Already:** Morphokinetic parameters obtained with time-lapse devices are increasingly used for human embryo quality assessment. However, their annotation is time-consuming and can be operator-dependent, highlighting the need of developing automated approaches.

**Study Design, Size, Duration:** This monocentric pilot study was conducted using 701 blastocysts originating from 584 couples undergoing IVF with embryo culture in a time-lapse device and on 4 mouse embryos.

**Participants/Materials, Setting, Methods:** An automated annotation tool was developed based on grey level coefficient of variation and detection of the thickness of the zona pellucida. The timings of cellular events obtained with the automated tool were compared with those obtained manually by 2 expert embryologists. The same procedure was applied on 4 mouse preimplantation embryos obtained with a different device in a different setting.

**Main Results and the Role of Chance:** Although some differences were found when embryos were considered individually, we found an overall excellent concordance between automated and manual annotation of human embryo morphokinetics from fertilization to expanded blastocyst stage (r^2^=0.94). Moreover, the automated annotation tool gave promising results across species (human, mice).

**Limitations, Reasons for Caution:** These results should undergo multi-centric external evaluation in order to test the overall performance of the annotation tool.

**Wider Implications of the Findings:** Our system performs significantly better than the ones reported in the literature and on a bigger cohort, paving the way for high-throughput analysis of multicentric morphokinetic databases, providing new insights into the clinical value of morphokinetics as predictor of embryo quality and implantation.

**Study Funding/Competing Interest(s):** This study was partly funded by Finox Forward Grant 2016.

**Trial Registration Number:** NA

## Introduction

In vitro fertilization (IVF) has greatly improved over the last decade. However, success rates are still not optimal and often variable across centers and countries, with multiple pregnancy still being an issue (European IVF-monitoring Consortium (EIM), 2017). This situation is partially explained by the current limitations of embryo quality assessment methods. In fact, morphological assessment remains the most common method to evaluate embryo implantation potential and still suffers from a lack of predictive power despite the implementation of well-defined consensus guidelines (Alpha Scientists in Reproductive Medicine and ESHRE Special Interest Group of Embryology, 2011). Although embryo morphology on the day of transfer has been shown to be correlated with the outcome of the IVF cycle (Rhenman et al., 2015), it has also been demonstrated that this evaluation suffered from a lack of inter and intra observers’ reproducibility (Paternot et al., 2009) and was not correlated to embryo ploidy (Capalbo et al., 2014).

Promoting single embryo transfer implies that accurate and relevant strategies to identify embryos with the best chance to implant are available (Kushnir et al., 2017). Various technologies, either invasive or non-invasive, have been developed over the last decade in order to improve embryo quality assessment in vitro, such as preimplantation genetic screening (Gardner et al., 2015), or metabolomics (Sanchez et al., 2017) with encouraging results, but also with some limitations such as cost, regulatory constraints and lack of clinical validation.

Time Lapse Monitoring (TLM) systems allow continuous and dynamic annotation of individual embryo development while maintaining optimal culture conditions (Basile et al., 2015; Castelló et al., 2016), bringing opportunities for personalized medicine approaches in IVF. The data generated are called morphokinetic parameters, as they combine morphological features and kinetic evaluation of embryo development. Since the first commercial devices became available in 2010, this technology has been implemented in many IVF laboratories for routine embryo culture and selection. It was recently reported that 17% of the US IVF labs possessed at least one TLM system in 2017 (Dolinko et al., 2017). This promising technology aims at correlating morphological features and kinetic parameters such as the timing of cellular cleavages or intervals as described by Ciray et al. (2014) with embryo quality and outcomes. To date, there are eight randomized clinical trials (RCT) evaluated the relevance of using morphokinetic parameters to select the embryo with the best chance of giving pregnancy (Armstrong et al., 2018). Of these, the initial hierarchical model described by Meseguer et al., (2011) was compared to classical morphological selection in three RCTs (Rubio et al., 2014; Semra Kahraman et al., 2012; Siristatidis et al., 2015) whereas Goodman et al., (2016) used their own algorithm. A recent meta-analysis concluded that the use of time-lapse monitoring was associated with a significantly higher ongoing clinical pregnancy rate, significantly lower early pregnancy loss and a significantly increased live birth rate compared with morphological selection (Pribenszky et al., 2017), whereas another review gave conflicting- results (Chen et al., 2017). Additionally, a validated predictive model based on multicentric morphokinetic databases analysis has been reported (Petersen et al., 2016), however no universally applicable morphokinetic marker or model to predict embryo implantation has yet been identified.

Most TLM systems allow the individual monitoring of embryo development and its manual annotation and record. Thus, trained operators have to regularly annotate morphokinetic parameters for each embryo in culture, which can be time-consuming depending on the number of cultured embryos and on the number of morphokinetic parameters annotated for each embryo. Moreover, the manual annotation of embryo developmental events can be impacted by inter-operator and inter-laboratory variability (Chen et al., 2013; Martínez-Granados et al., 2017; Sundvall et al., 2013), preventing robust and high throughput multicentric databases analysis. However, embryo annotations through TLM systems continues to be performed manually and suffers from the same inter-operator variability as microscopic observation (Martínez-Granados et al., 2017). Moreover, the type of TLM system can contribute to increasing embryo annotation discrepancies. Automated annotation of embryo morphokinetic parameters could theoretically tackle these 2 issues, leading to improved workflow, high throughput analysis, and lower subjectivity. However, a commercially available automated TLM system based on few early parameters recently failed in confirming its clinical relevance in a prospective trial (Kieslinger et al., 2016), highlighting the need for a robust and multiparametric automated annotation tool. To our knowledge, only one pilot study reported the set-up of a semi-automated script annotating embryo morphokinetic parameters in humans (Mölder et al., 2015). This script was, however, tested on a very limited population (39 embryos) and was limited to a proof of concept. In the study described here, we have greatly improved the method and enhanced its robustness and detection power.

The purpose of this study was to test the performance of a new fully automated tool called “Kinetembryo” for the annotation of embryo morphokinetic parameters obtained with a commercially available time-lapse incubation device on human embryos. Secondly, we evaluated the robustness of the tool by performing a cross-validation pilot study on mouse embryos.

## Materials & Methods

In order to evaluate the performance of our method and software tool Kinetembryo for the fully automatic annotation of embryo morphokinetic parameters, we compared the annotation performed manually as detailed below with the automatic output of Kinetembryo on the same movies.

### Design

This monocentric study was conducted in a University-based fertility center with data from unselected couples referred for ICSI and whose embryos were cultured in the Embryoscope^®^ (Vitrolife, Sweden) between 16/02/2011 and 01/01/2017. All patients gave their consent for anonymous use of their data registered in this database. This project was approved by the local Institutional Review Board (GNEDS). All patients underwent controlled ovarian stimulation with antagonist protocol as described in a previous study (Fréour et al., 2015).

### Embryology Procedures

ICSI and embryo culture were performed as described previously (Fréour et al., 2015) at 37°C under controlled atmosphere with low oxygen pressure (5% O2, 6% CO2). Sequential media were used for embryo culture (G1plus^®^ and G2plus^®^, Vitrolife). Images were captured on seven focal plans every 15-min intervals using Hoffman modulation contrast (HMC) optical setup (Hoffman, 1977) and a 635 nm LED as light source as provided in the Embryoscope^®^. The resolution of the camera is 1280×1024 pixels. In this study 704 movies were used for the performance evaluation of the method, all of them having been annotated as a reference.

The development of each embryo was prospectively annotated as described by Ciray et al., (2014) by two trained embryologists undergoing regular internal quality control (four times per year) in order to keep inter-operator variability as low as possible. Zona Pellucida (ZP) thickness was measured at two opposite points. The terms t2, t3, t4, t5, t6, t7, t8 and t9+ were respectively used for exact timings of blastomere cleavage resulting in 2, 3, 4, 5, 6, 7, 8 and 9 well-defined blastomeres. The term tM referred to a fully compacted morula. At the blastocyst stage, tSB was used to describe the onset of a cavity formation, tB was used for full blastocyst i.e. the last frame before the Zona Pellucida (ZP) starts to thin, tEB for expanded blastocyst, i.e. when the ZP is 50% thinned. Blastocyst contractions and the beginning of herniation were also recorded.

### Development of an Automated Annotation Tool

The development procedure consisted of 2 main steps: a preprocessing step and an automatic annotation step (Figure 1). It was implemented using stand-alone software written with Matlab^®^. The input was the movies generated by the TLM system, in our case the Embryoscope^®^, which are 2D movies of the optimal in-focus plan of acquisition.

**Figure 1:**
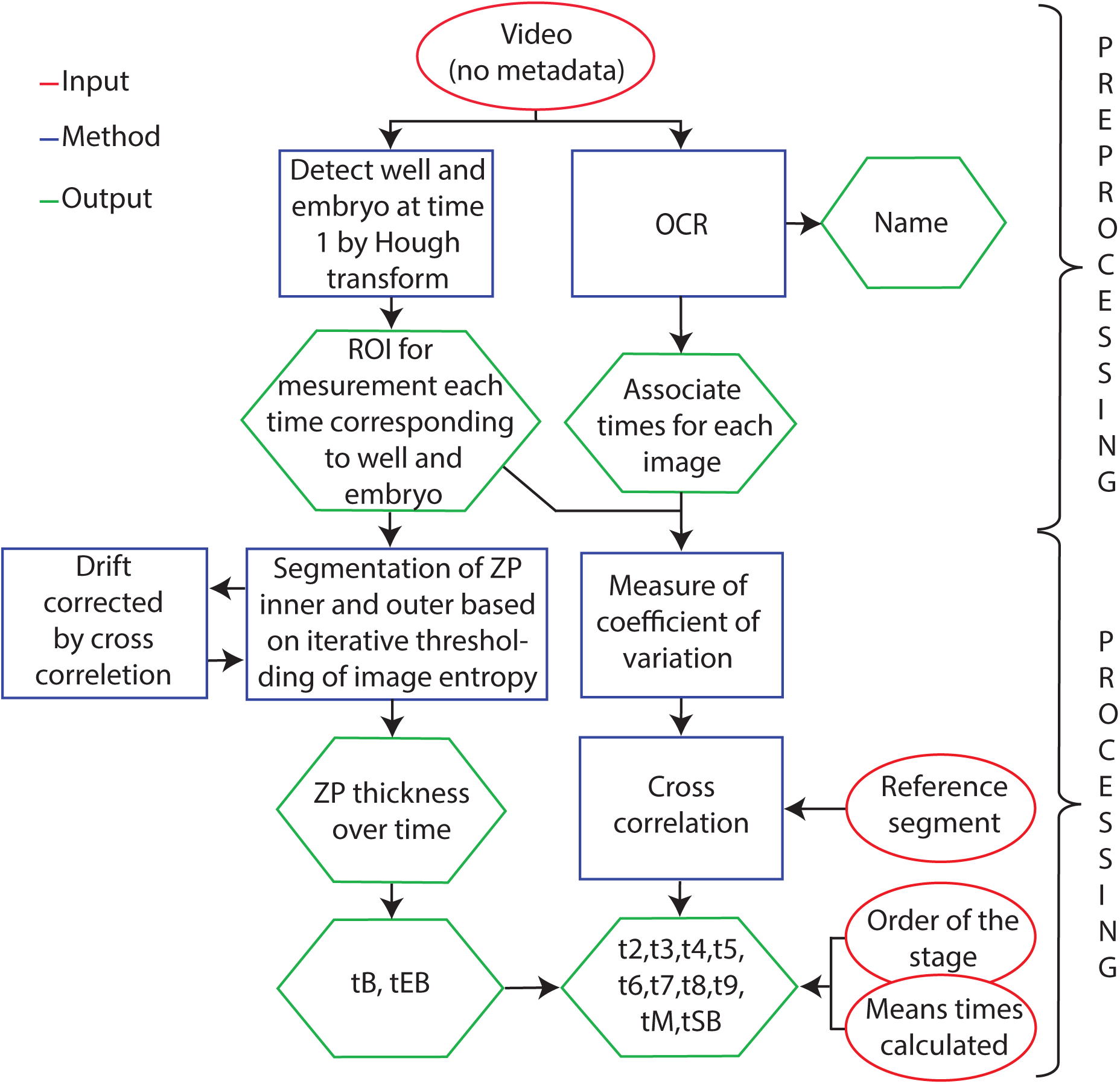
Schematic Representation of the Process Used to Develop the Automated Annotation Tool.

First, the preprocessing step consisted in extracting metadata, the name and times, and region of interest (ROIs), from the well and the embryo cell.

Then, the annotation step consisted of two main tasks. The first task allowed detection of tB and tEB. The Zona Pellucida (ZP) (inner and outer) was segmented. Time points, tB and tEB, were then automatically detected from the ZP thickness curve (Figure 2). The second task consisted in identifying earlier events t1 to t9, tM and tSB. For this, the approach used was inspired from the study of Mölder et al., (2015). In our work, we measured the grey-level coefficient of variation of intensity into the tracked embryo ROI, giving a curve of variation over time (Figure 3). To detect the different key stage of embryo development from this curve, first we developed the fragment curve as a reference, based on the observation of curves. For each reference fragment, the temporal location of the most similar part of the curve is searched for by cross-correlation. This narrowed down our search area for stages.

**Figure 2:**
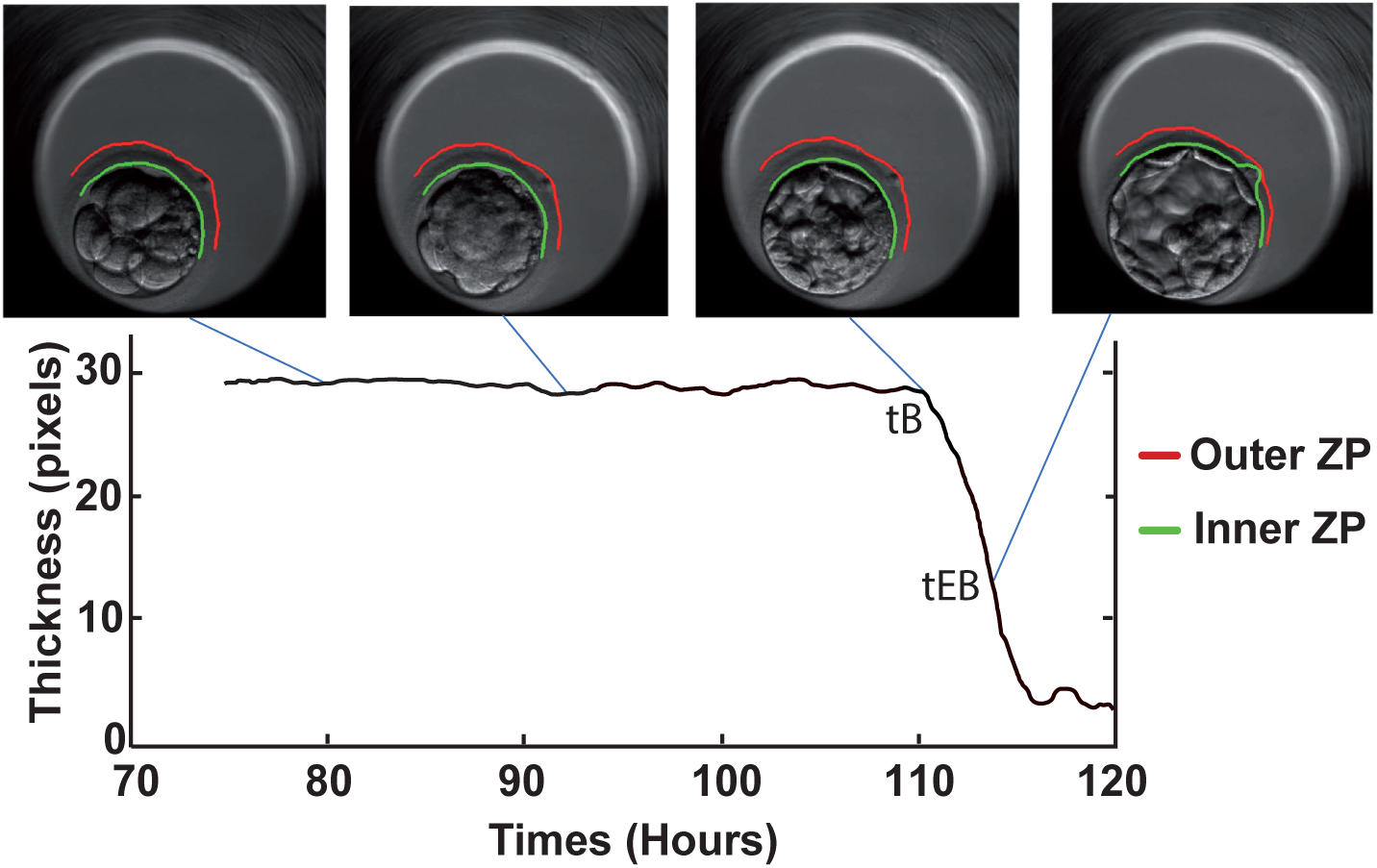
Representative Illustration of Zona Pellucida Thickness Measurement.

**Figure 3:**
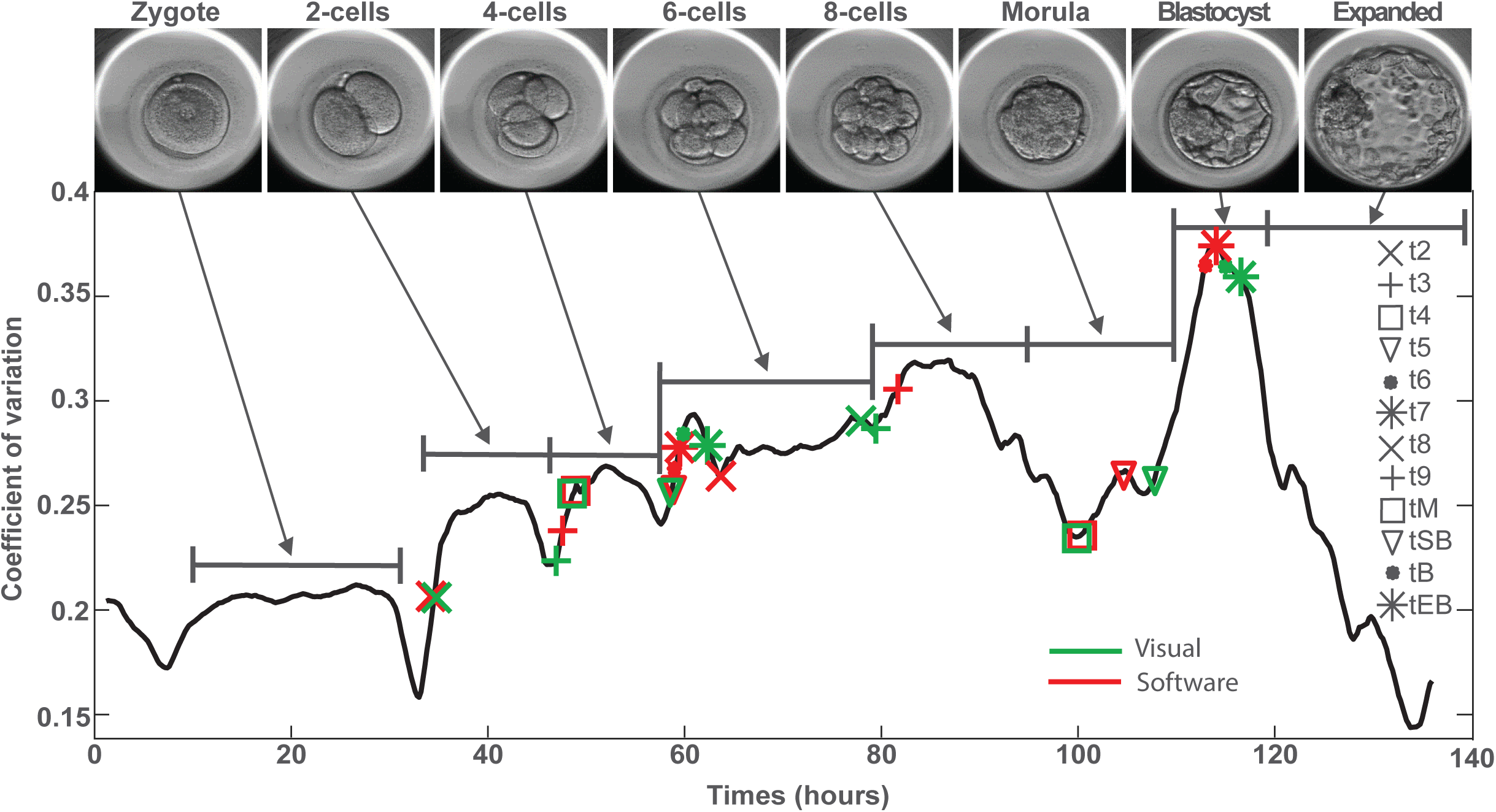
Representative illustration of the evolution of grey level coefficient of variation throughout embryo development from fertilization to expanded blastocyst stage.

### Validation of the automated annotation tool

In order to test the performance and the accuracy of this automated analysis process, we compared the results with the timings prospectively recorded by 2 expert users on a large set of 701 individual videos extracted from the Embryoscope^®^, corresponding to embryos monitored up to the blastocyst stage and used for blastocyst transfer. Regression analysis was used to evaluate the association between the expert annotation and the automatic annotation process. We further tested the hypothesis of identical mean results for the detection of each embryo stage between both methods by Student’s t-test. A difference <2 hours between both methods was arbitrarily considered as acceptable for the annotation of each embryo developmental step.

In order to test the flexibility of Kinetembryo on other species and other imaging system, the software was ran on movies generated by 4 confocal time-lapse records of mouse preimplantation embryos cultured as described by Raveux et al. (2017). Embryos were imaged using a Zeiss LSM800 equipped with a Plan-apochromat 20X/NA 0.8 objective on multiples focal plans every 10 or 30 minutes intervals. Movies were manually annotated by an expert following the same nomenclature as for human embryos. For the automatic analysis, the same reference curve computed for human embryo analysis was used. The only difference in the processing was to skip the ZP detection and to get the metadata directly from the movies. The a priori knowledge was also adapted (literature times, and order of morula event).

## RESULTS

### Comparison of automated vs manual grading of preimplantation embryos

To test the robustness of Kinetembryo, data from 701 movies of embryo development originating from the Embryoscope with full follow-up from fertilization up to blastocyst stage was applied. For each movie, the time associated with the beginning of a stage obtained by automated or manual annotation was used (Figure 4). When comparing both annotation methods, an overall significant correlation with a R^2^ value of 0.94 was found, supporting the high degree of sensitivity of Kinetembryo tool.

**Figure 4:**
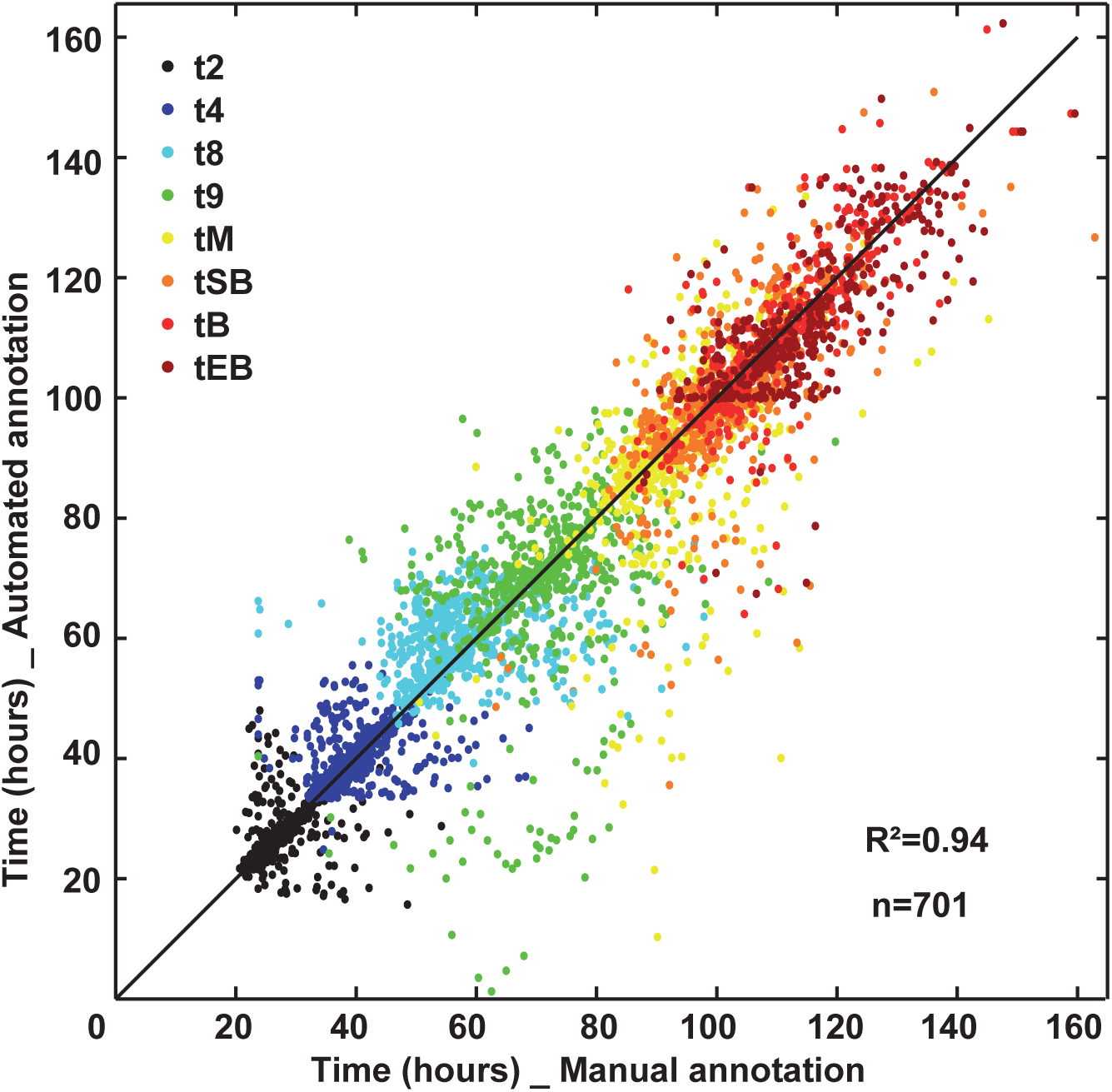
Performance test of the automated annotation tool, as reflected by correlation analysis between automated and expert manual morphokinetic annotation of 701 full embryo development monitored with time-lapse device.

Time of processing on a personal computer was on average 20 minutes and did not require any manual intervention, allowing to batch 701 movies together. Note that the code have been parallelized on the data, meaning that when ran on several processors, several movies were processed in parallel, reducing the total time of processing.

### Mitotic divisions

Kinetembryo was able to correctly annotate most of the first mitotic divisions from 2-cell stage and up to 6-cell stage within a 2 hours range when compared with manual annotation (Figure 5). The following stages were correctly annotated for less than half of the samples (42.5% for t7, 28.9% for t8, 27.8% for t9). At the end, mean cleavage timings given by automated annotation were not statistically different from those given by manual annotation from t2 to t6. However, the difference between the 2 methods became statistically significant from t7 to t9 (p < 0.05).

**Figure 5:**
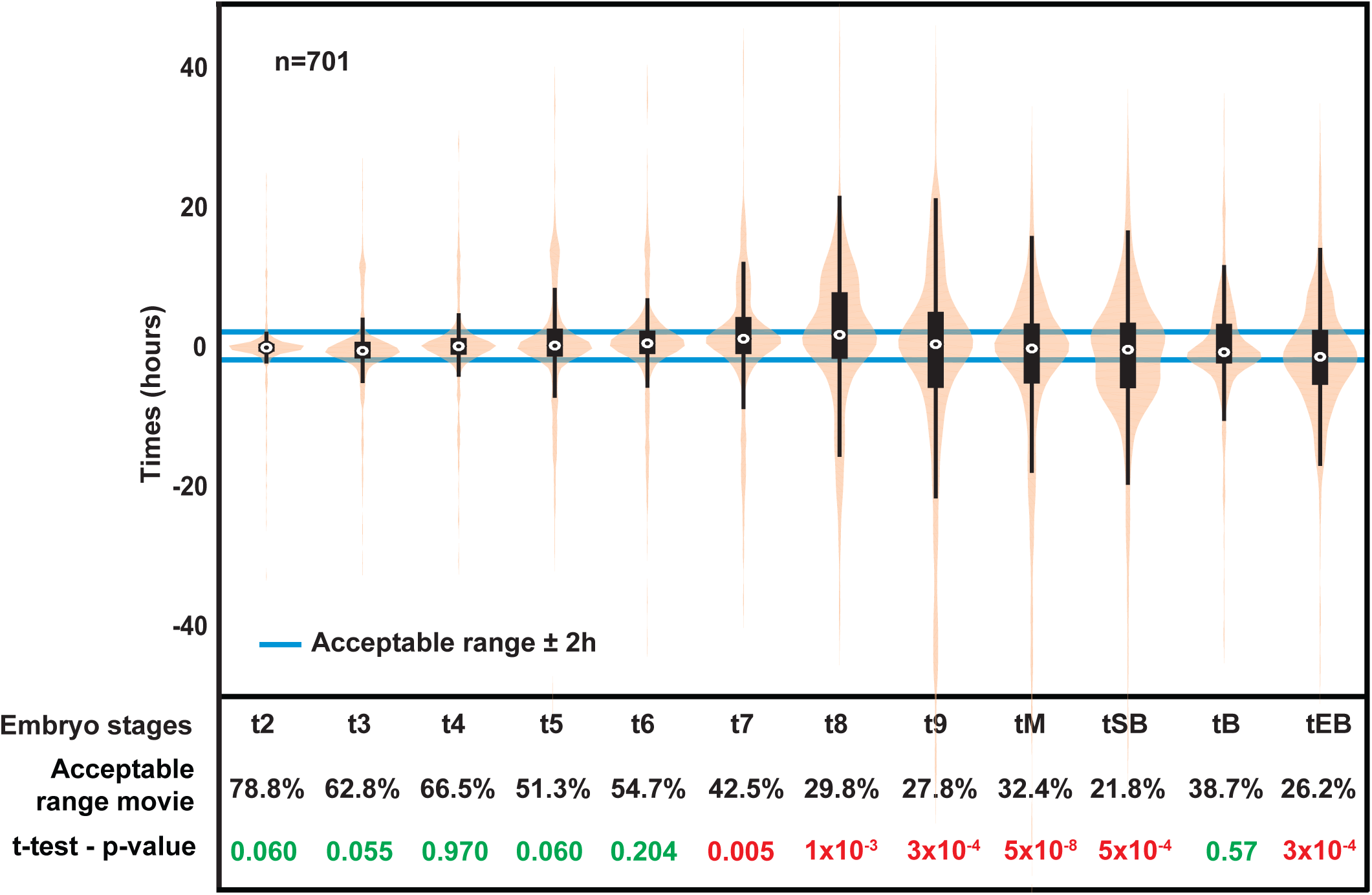
Performance test of the automated annotation tool, as reflected by the graphical representation (boxplot: in black and the point for the average) and the density (violin plot: in beige) of the difference between manual annotation and automatic each embryonic stage.

### Compaction and blastulation

The full compacted morula stage was well detected by Kinetembryo in 32.4 % of the samples. The following steps of blastulation (tSB and tEB) were not accurately annotated by Kinetembryo with respectively 21.8 and 26.2 % of annotation within the range of acceptable error (<2 hours) against manual assessment but tB could be correctly annotated for 38.7 % of the embryos. Nevertheless, tB stage annotation was not statistically different between manual and automated annotation (p = 0.57).

### Cross-validation of our automated process on animal dataset

To adapt to the growing number of TLM systems available on the market, a software should be able to work on multiple platforms. To demonstrate the robustness and adaptability of our system, we applied our automated process to 4 time-lapse records of mouse preimplantation embryos obtained with a different device in a different setting. The curve of variation of grey in the mouse video has a variation similar to the human video (Figure 6). Moreover, Kinetembryo was able to detect a mouse specific event: early compaction, which is distinguished by a step of compaction and occurs at the 8 cells stage.

**Figure 6:**
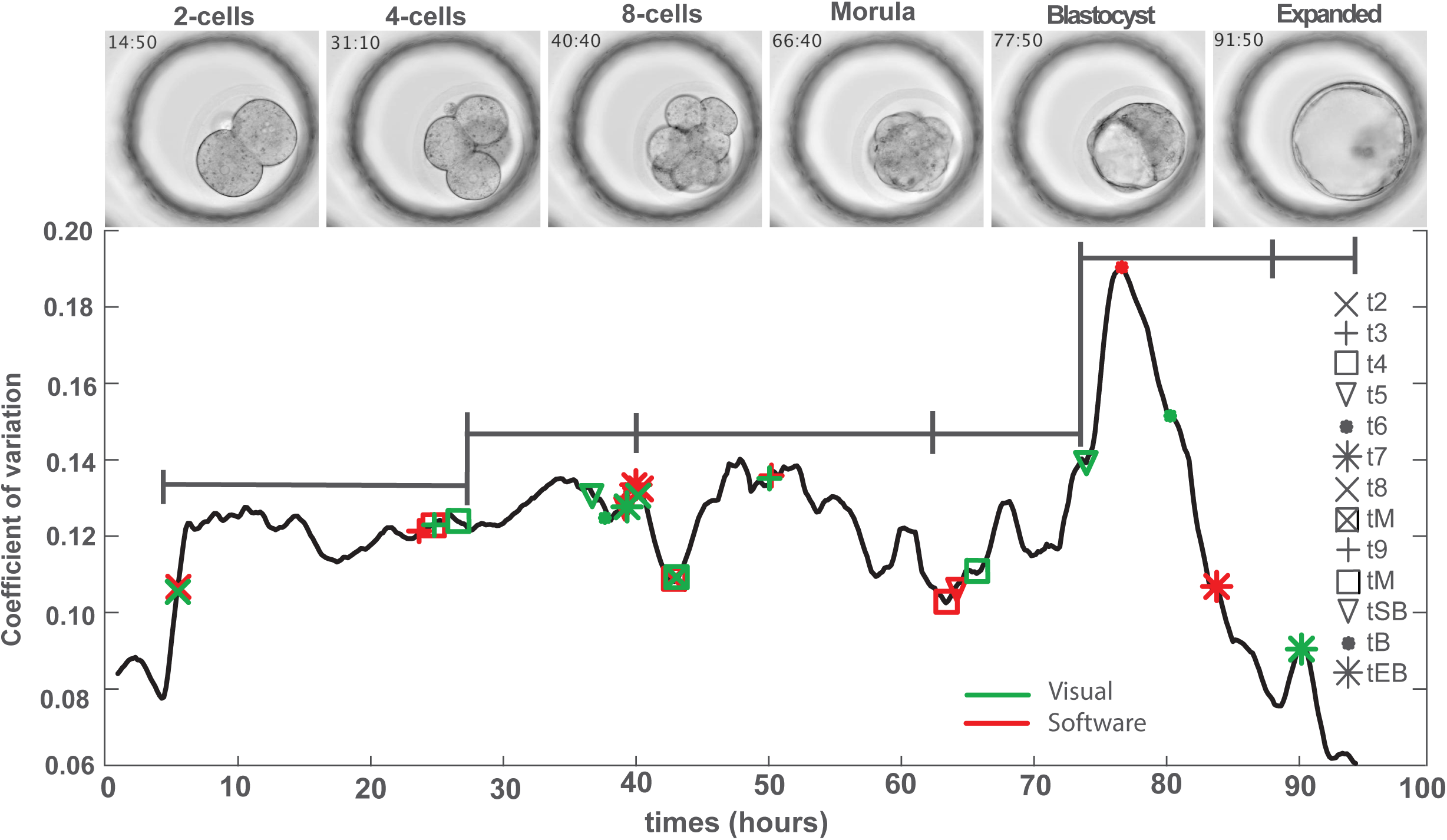
Performance test of the automated annotation tool on videos of mouse embryo development. Representative illustration of the evolution of grey level coefficient of variation throughout embryo development from fertilization to expanded blastocyst stage.

## DISCUSSION

In this study, we showed that our automated tool Kinetembryo offered a reliable tool for quick annotation of embryo morphokinetic parameters. Although some differences were found when embryos were considered individually, this tool could help to set up high-throughput studies in large multicentric databases in a reproducible way. Moreover, the possibility to automatically annotate mouse embryo development as well paves the way for inter-species morphokinetic events comparison.

Time Lapse Monitoring (TLM) of embryo development applied to assisted reproductive technologies (ART) raises hope of better clinical outcome for infertile couples thanks to optimal embryo culture conditions and objective criteria for embryo selection in a single embryo transfer strategy. However, its clinical effectiveness over conventional morphology is still under discussion. In a recent meta-analysis, Chen et al. (2017) showed that clinical TLM may have the potential to improve outcomes but that more evidence was needed, while Pribenszky et al. (2017) concluded in another meta-analysis to an improvement of pregnancy rates and lower pregnancy losses when using TLM. The pros and cons were recently debated in a review on the main topics related to TLM (Paulson et al., 2018). Overall, experts in both groups provided sound but opposing arguments, and finally reached the conclusion that robust and appropriately designed clinical trials are needed to draw a firm conclusion on the clinical usefulness of TLM in ART. It should be noted that financial aspects remain critical as well when discussing about TLM.

Such multicentric RCTs in large cohorts imply the analysis of huge databases originating from various settings. This implies time-consuming manual annotation of embryo morphokinetic parameters in each participating center and large data management capacity. Moreover, the analysis of such results implies robust inter-centric embryo annotation, limiting as much as possible inter operator variability (Martínez-Granados et al., 2017; Sundvall et al., 2013). Automated analysis and annotation of early embryo development represents the best way to reach such high standards. In this study, we have shown that our automated image analysis process, called Kinetembryo, was able to proceed to a full embryo annotation from first cleavage up to expanded blastocyst stage. Moreover, we observed excellent correlation with manual assessment validated on a large study population, with a better accuracy for early stages than for late embryo stages. Up to now, most of the published embryo selection algorithms using morphokinetic events were mainly based on early morphokinetic events such as first cleavages (Petersen et al., 2016; Rubio et al., 2014). Following this strategy, Eeva^®^, a semi-automated annotation tool for early embryo events detection was developed with the aim of improving the selection of embryos with high developmental potential for early stages embryo transfer in addition with day 3 morphology (Conaghan et al., 2013). The use of Eeva^®^ in addition with standard morphology assessment has been compared with standard morphology assessment in few randomized clinical trials with different conclusions. On one hand Eeva^®^ seemed to give objective information for helping embryologists with various levels of experience to choose the best embryo to transfer in a double-blinded multi-center study with 54 patients undergoing blastocyst transfer (Diamond et al., 2015), on the other hand, an observational prospective two-center study did not demonstrate any improvement in cycle outcome when using Eeva^®^ (Kieslinger et al., 2016).

The automated process described here whilst providing very promising results should be further refined in order to improve the detection of later stages of embryo preimplantation development. The next step would be to undergo external validation in a multicentric design, regardless of the commercial TLM device used in the participating IVF centers. Once validated, this kind of automated process could solve the main drawbacks of most of the existing studies comparing TLM and morphological embryo assessment which are a lack of robustness, objectivity and statistical power due to monocentric analysis (Armstrong et al., 2018). This cross validation between centers is necessary as limitations of predictive models have been shown when tested externally (Barrie et al., 2017; Fréour et al., 2015). Moreover, Storr et al. (2018) recently described discrepancies between published selection models and morphological assessment selection of the embryo to transfer by trained embryologists.

These results suggest that Kinetembryo could move the field closer to better TLM standards, improving ART outcomes. The next improvement will be to design a process which automatically grades embryos. With such an improvement, ART could achieve another breakthrough leading to improved results.

## CONCLUSION

Overall, a robust automated morphokinetic annotation process has been developed, which can easily be transferred to multiple platforms and species. This tool could pave the way to a refined, deeper understanding of human preimplantation development, and support further discovery of predictive tests to assist embryologists for embryo selection ultimately improving ART success rates.

## Author’s roles

MF: drafted the manuscript, development of the automated annotation tool, analysis of the results

AR: drafted the manuscript, manual time-lapse annotation, analysis of the results

MM: development of the automated annotation tool

JL: manual time-lapse annotation

DM: expert knowledge in bioinformatics, critical revision of the manuscript

SVP, MCT: generated and provided mouse videos

PB: critical revision of the manuscript

PPG: study design, supervision of the study, expert knowledge in image analysis

LD: study design, supervision of the study, critical revision of the manuscript

TF: study design, supervision of the study, critical revision of the manuscript, validated the final version of the manuscript

## Funding

This study was partly funded by Finox forward grant 2016

